# Optimizing brinjal (*Solanum melongena* L.) health and yield through bio-organic amendments against Fusarium wilt

**DOI:** 10.1101/2025.05.13.653889

**Authors:** Jannatul Mauya, Khadija Akhter, Hasiba Kabir Mim, Marzia Rahman, Raihan Ferdous

**Affiliations:** Department of Plant Pathology, Sher-e-Bangla Agricultural University, Dhaka-1207, Bangladesh; Mushroom Development Institute, Savar, Dhaka-1340, Bangladesh

**Keywords:** Biochar, Disease incidence, Eggplant, Spent mushroom substrate, Yield

## Abstract

A field experiment was conducted to evaluate the effect of different bio-organic treatments, including control (T_0_), spent mushroom substrate-SMS (T_1_), vermicompost (T_2_), poultry manure (T_3_), biochar (T_4_), SMS with biochar (T_5_), SMS with poultry manure (T_6_), and SMS with vermicompost (T_7_) on the management of Fusarium wilt of brinjal (*Solanum melongena* L.). Pathogen identification confirmed *Fusarium oxysporum* f. sp. *melongenae* as the causal agent of wilt through morphological and pathogenicity tests. Growth attributes, yield, and disease incidence were recorded at various growth stages. The lowest disease incidence (5.55%) was observed in T_5_, followed by T_6_, while the highest disease incidence (44.4%) was found in the untreated control (T_0_). The treatment of T_5_ significantly enhanced plant growth parameters, including plant height, number of leaves, and number of branches, alongside a substantial increase in yield (12.71 tons/ha), which was statistically similar to T_6_. These bio-organic amendments not only suppressed disease incidence effectively but also improved soil health, enhanced microbial diversity, and promoted vegetative growth and yield. The results indicate that the integration of SMS with Biochar or Poultry Manure is a sustainable and eco-friendly strategy for managing Fusarium wilt in brinjal cultivation, potentially replacing conventional chemical methods for enhanced productivity and soil health.

## Introduction

Brinjal (*Solanum melongena* L.) is the most popular, widely available, and reasonably priced vegetable that belongs to the family Solanaceae. In Bangladesh, “Begoon” (also known as brinjal or eggplant) is a common and beloved vegetable that has a connection to the social, cultural, and rural residents’ financial circumstances. In the Kharif (summer) season in Bangladesh, 49406.92 acres were cultivated, with total production at 209541.17 MT, and in the Rabi (winter) season, 84536.65 acres were cultivated, with total production 409001.46 MT [1]. The fruit of the brinjal is very low in calories and has a healthy amount of minerals. Brinjals are rich in anthocyanin chemicals. The strong protective effects of anthocyanin against diabetes, cancer, cardio vascular disease. [2]. Diseases caused by fungi, bacteria, viruses, and nematodes severely reduce the amount of brinjal produced [3]. Wilt of brinjal is one of the most harmful diseases among all the diseases of brinjal that causes great economic loss of brinjal production. Fusarium wilt initiated by *Fusarium oxysporum* f. sp. *melongenae* are the most devastating one. Both the quality and yield of the brinjal production are decreased by fusarium wilt. The severity of fusarium wilt in brinjal is 10%-90% [4]. Fusarium wilting generally causes 20–30% of eggplant plants to die and it may turn into epidemic during November to December [5]. *Fusarium oxysporum* can survive in the soil for a long period of time without any host. Because *Fusarium oxysporum* is a soil-borne pathogen that makes it challenging to regulate. Several control measure such as crop rotation, the use of various resistant cultivars, soil sterilization and solarization and the application of fungicides are some useful or effective management strategies for Fusarium wilt. However, due to the possible negative impacts of fungicides on the environment and human health as well as their unfavorable effects on nontarget organisms, the extensive use of chemical fungicides has been a source of public worry and security [6]. Biological control could be successful alternative to chemicals. Several biocontrol agents such as spent mushroom substrate, poultry refuse, vermicompost have the ability to manage some soil borne pathogen. Because of its high cation exchange capacity (CEC), slow mineralization rate, and rich nutritional status, the SMC has been discovered to be an excellent source of nutrients for agriculture [7]. SMS is rich in diverse microorganisms, such as disease antagonistic bacteria and fungus. It is biodegradable, safe to apply and less expensive to develop [8]. Vermicomposting is the natural process of rotting or decomposition of organic matter by the activity of earthworm under controlled conditions [9]. Vermicompost’s organic carbon slows the release of nutrients into the system so that they can be absorbed by the plant. [10]. Poultry manure is an effective soil amendment that supplies nutrients for growing crops. Poultry manure has a high nutritional value since it contains nitrogen, phosphorus, and potassium. [11]. The incidence of wilt (*F. oxysporum*) can be reduced 41.15% by poultry manure [12]. Biochar is a type of charcoal that is produced through a process called pyrolysis, which involves heating organic materials (such as wood chips, agricultural waste, or animal manure) in the absence of oxygen. Biochar has been utilized to boost agricultural productivity, enhance soil health, and lower greenhouse gas emissions [13]. This study proposes the use of spent mushroom substrate to combat the pathogen *Fusarium oxysporum* and to find out the effectiveness of Spent Mushroom Substrate and other soil amendments for the management of wilt of brinjal (*Fusarium oxysporum*).

## Materials and methods

A research experiment was conducted to evaluate the effect of spent mushroom substrate and other organic amendment against fungal wilt of brinjal (*Fusarium oxisporum* f. sp. *melongenae*).

### Study site and period

The field experiment was conducted in the Central Farm of Sher-e-Bangla Agricultural University and lab experiment was conducted in MS laboratory of the Department of Plant Pathology, Sher-e-Bangla Agricultural University, Dhaka-1207. The experiment was carried out during the period from September 2021 to April 2022.

### Selection of varieties and treatments

Brinjal variety BARI BT Brinjal 2 (Kajla) was used for the experiment and the variety was collected from Bangladesh Agricultural Development Corporation (BADC), Gabtoli, Dhaka. Along with control, seven treatments were selected and evaluated through a field experiment in natural conditions (*in vivo*). The treatments were used in this study are following:

T_0_ = Control,

T_1_ = Spent Mushroom Substrate (SMS),

T_2_ = Vermicompost,

T_3_ = Poultry manure,

T_4_ = Biochar,

T_5_ = Spent Mushroom Substrate + Biochar,

T_6_ = Spent Mushroom Substrate + Poultry manure and

T_7_ = Spent Mushroom Substrate + Vermicompost.

The chosen treatments were applied to the main field 20 days before transplanting the seedlings to ensure adequate decomposition, the growth of competitive microorganisms, and the development of the pathogen-suppressing abilities.

### Experimental design

The design of the study was laid out in Randomized Complete Block Design (RCBD) with eight treatments and three replications. The experimental field was divided into three blocks. Each blocks had eight plots. As a result, three blocks had 24 plots. The size of each plot was (2.5 × 1.8) m. The space was 0.75 m kept between the blocks and 0.50 m was kept between the plots. Each plot had 6 plants and plant to plant distance was 75 cm.

### Data collection

On the basis of observable symptoms of fusarium wilt of brinjal, responses to the selected different treatments were noted at 25, 45, and 65 DAT (Days After Transplanting). The following parameters were considered for data collection-percent disease incidence, plant height, number of branches per plant, number of leaves per plant, number of fruits per plant, number of fruits per plot, fruit length, individual weight of fruits, yield per plant, yield per plot, yield per ha. Yield per plant and disease incidence were calculated following the formula [14]:

Where,

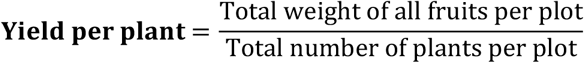

And

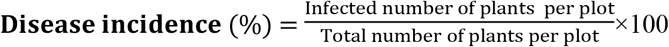

### Isolation and identification of the pathogen

Infected stem was collected from the experimental field, and the stem was cut into small pieces (.05-1cm) by a sterilized knife. Sterilization was done by dipping in 0.01% mercuric chloride (HgCl_2_) for 2–3 minutes, and then placed on blotter paper and then incubated at 25±1°C for 7–10 days. When the fungus was grown then transferred to PDA media. The pathogen was then purified by the transfer of mycelium from the tip of the colony [15]. Finally, a compound microscope was used to study the morphological characteristics of the fungus.

### Pathogenicity test under pot culture

The soil was taken from the field. After that, 0.4% formalin solution was completely mixed with soil @ 200ml/cft soil and kept under a polythene sheet for 48 hours to retain the gases within the soil for sterilization. The soil was then exposed to sunlight for seven days. After 7 days, the treated soil was ready to use. The soil was then placed in 25 cm diameter surface-sterilized pots. BT Brinjal 2 seedlings were raised in a plastic pot. Sterilized soil having fertilizers as per the package of practices was used for seedbed preparation. Thirty-day-old seedlings of brinjal were treated with spore suspension of *F. oxysporum* by the root dip method. Damaged roots were immersed in a conidial suspension (10^6^ conidia/ml) for 10 minutes while the control plants were plunged in sterile tap water. After that, seedlings were placed into clean pots. The plants were watered regularly and observed for the appearance of wilt symptoms. Observations were done on wilt symptoms for up to 5 weeks. After three weeks of inoculation, symptoms were seen on the inoculated plant, and the pathogen was reisolated and compared with the original culture of *F. oxysporum* to satisfy Koch’s postulates [15].

### Statistical analysis

The data were statistically analyzed by using the computer-based software Statistix 10 and by using analysis of variance (ANOVA) to find out the variation of results from experimental treatments. Treatment means were compared by LSD.

## Results

### Study on pathogen identification

The pathogen *Fusarium oxysporum* f. sp. *melongenae* was identified through microscopic examination based on symptomatology. On Potato Dextrose Agar (PDA) media, the fungus produced whitish to light pink mycelium, which gradually expanded into a light gray colony during sporulation. Microscopic analysis of the pure culture revealed the presence of 2–3 celled, slightly curved macroconidia, alongside single-celled microconidia (Fig. 1).

**Fig. 1.**
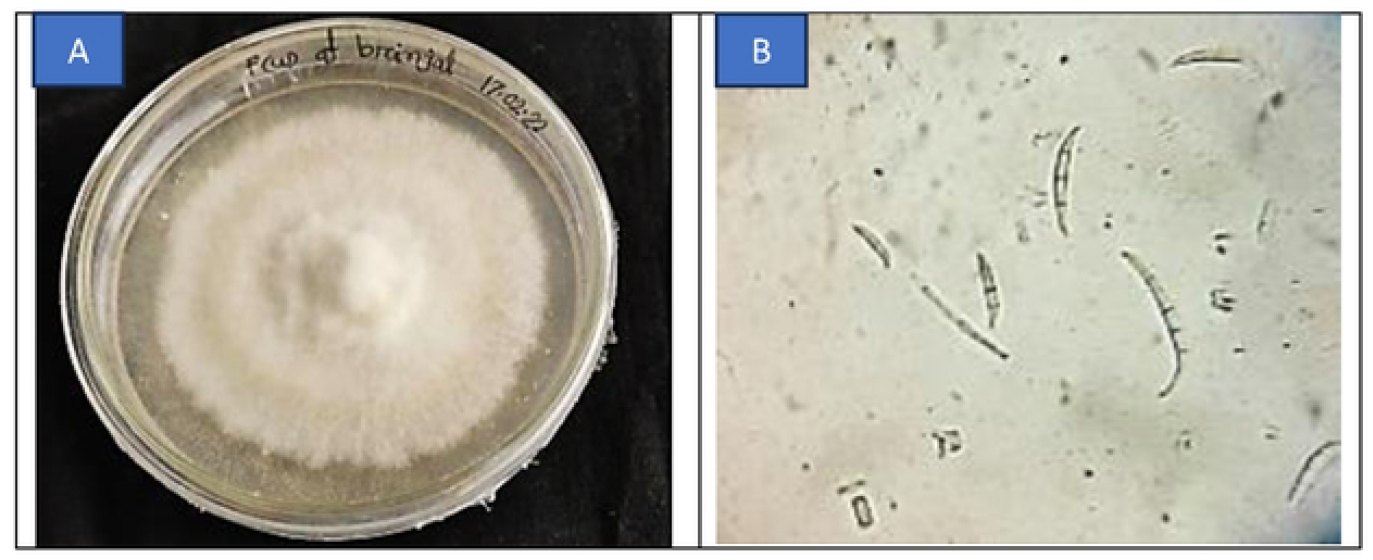
Morphological study of *Fusarium oxysporum* f. sp. *melongenae* **A.** Pure culture on PDA media, and **B.** Microscopic view under a compound microscope (40X)

Pathogenicity was confirmed through typical wilting symptoms observed in brinjal plants inoculated with the isolated fungus. Re-isolation of *Fusarium oxysporum* f. sp. *melongenae* from the stems of inoculated, wilt-affected plants completed Koch’s postulates, verifying its role as the causal agent of brinjal wilt.

### Study on disease incidence

The influence of the selected bio-organic treatments on disease incidence was evaluated at 25, 45, and 65 Days After Transplanting (DAT). At 25 DAT, the untreated control (T_0_) exhibited the highest disease incidence (44.4%), whereas the lowest incidence (5.55%) was recorded in T_5_ (Spent Mushroom Substrate + Biochar). Similar trends were observed at 45 DAT and 65 DAT, where T_0_ maintained the highest disease incidence (55.55% and 77.77%, respectively), while T_5_ continued to show the most effective disease suppression (16.66% at both intervals).

Field observations confirmed that T_5_ effectively controlled disease incidence across all time points, followed by T_6_ (Spent Mushroom Substrate + Poultry Manure), highlighting the potent suppressive effect of organic amendments like spent mushroom substrate and biochar (Table 1).

**Table 1.** Impact of different treatments on wilt incidence of brinjal at different Days After Transplanting (DAT)

| Treatments | Disease Incidence (%) at 25 DAT | Disease Incidence (%) at 45 DAT | Disease Incidence (%) at 65 DAT |
| --- | --- | --- | --- |
| T<sub>0</sub> | 44.44 a | 55.55 a | 77.77 a |
| T<sub>1</sub> | 16.66 b-d | 22.22 bc | 33.33 bc |
| T<sub>2</sub> | 16.66 b-d | 38.87 ab | 49.99 b |
| T<sub>3</sub> | 27.77 a-c | 33.33 bc | 49.99 b |
| T<sub>4</sub> | 16.66 b-d | 38.87 ab | 44.44 b |
| T<sub>5</sub> | 5.55 d | 16.66 c | 16.66 c |
| T<sub>6</sub> | 11.10 cd | 27.77 bc | 27.77 bc |
| T<sub>7</sub> | 33.33 ab | 38.87 ab | 44.44 b |
| CV (%) | 58.83 | 35.99 | 35.97 |

### Study on the growth parameters of brinjal under different treatments

The treatments significantly influenced plant growth parameters, including plant height, the number of leaves per plant, and the number of branches per plant. The highest plant height (71.0 cm) was achieved in T_5_, followed by T_6_ (66.33 cm), while the lowest (50.66 cm) was observed in T_0_ (Fig. 2). For the number of leaves per plant, T_5_ again exhibited the maximum count (111), significantly higher than T_0_ (73.67), indicating improved vegetative growth. Similarly, the highest number of branches per plant (11.33) was recorded in T_5_, statistically similar to T_6_ (10.66), whereas the lowest (5.67) was found in T_0_ (Fig. 3).

**Fig. 2.**
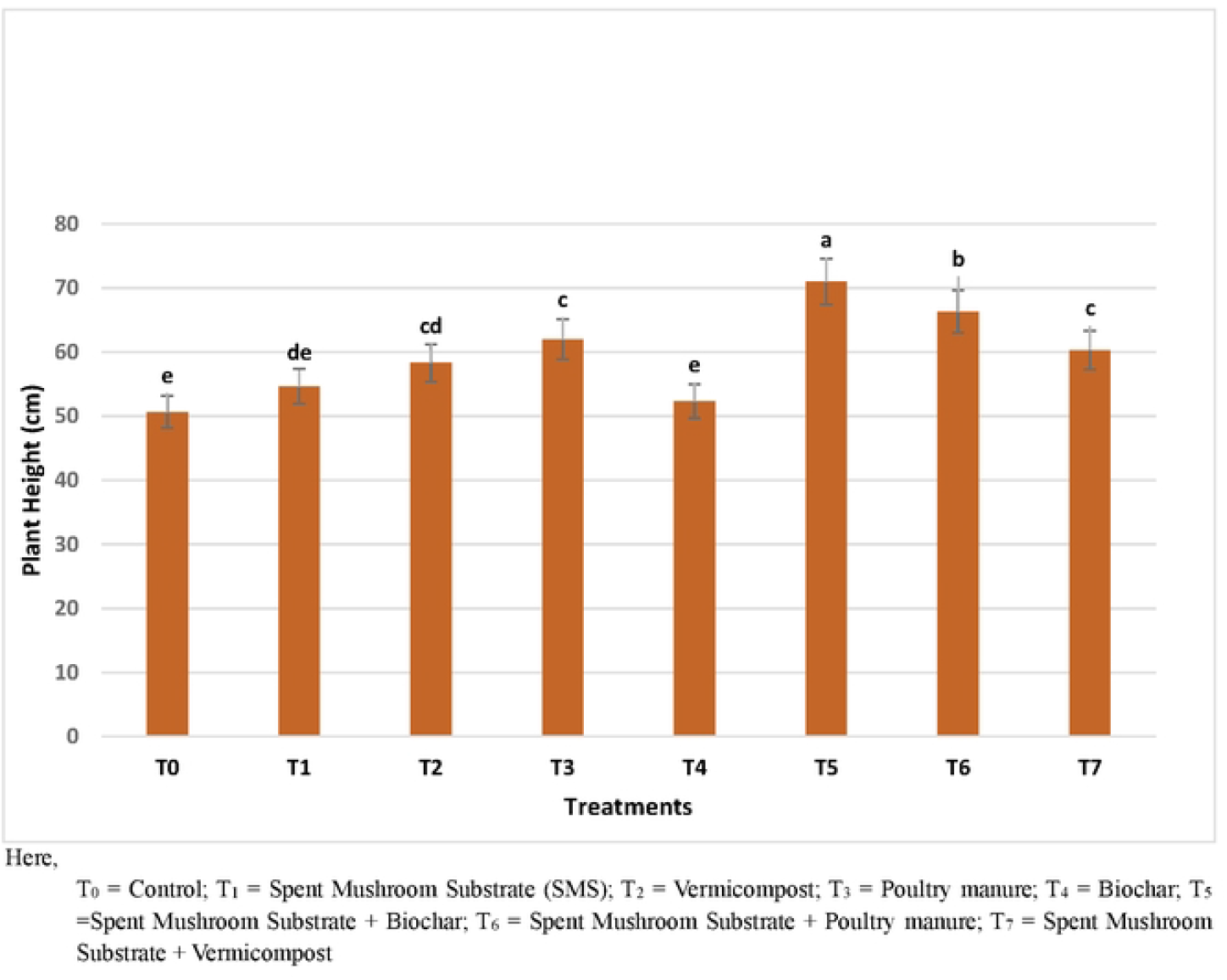
Impact of different treatments on the plant height of brinjal

**Fig. 3.**
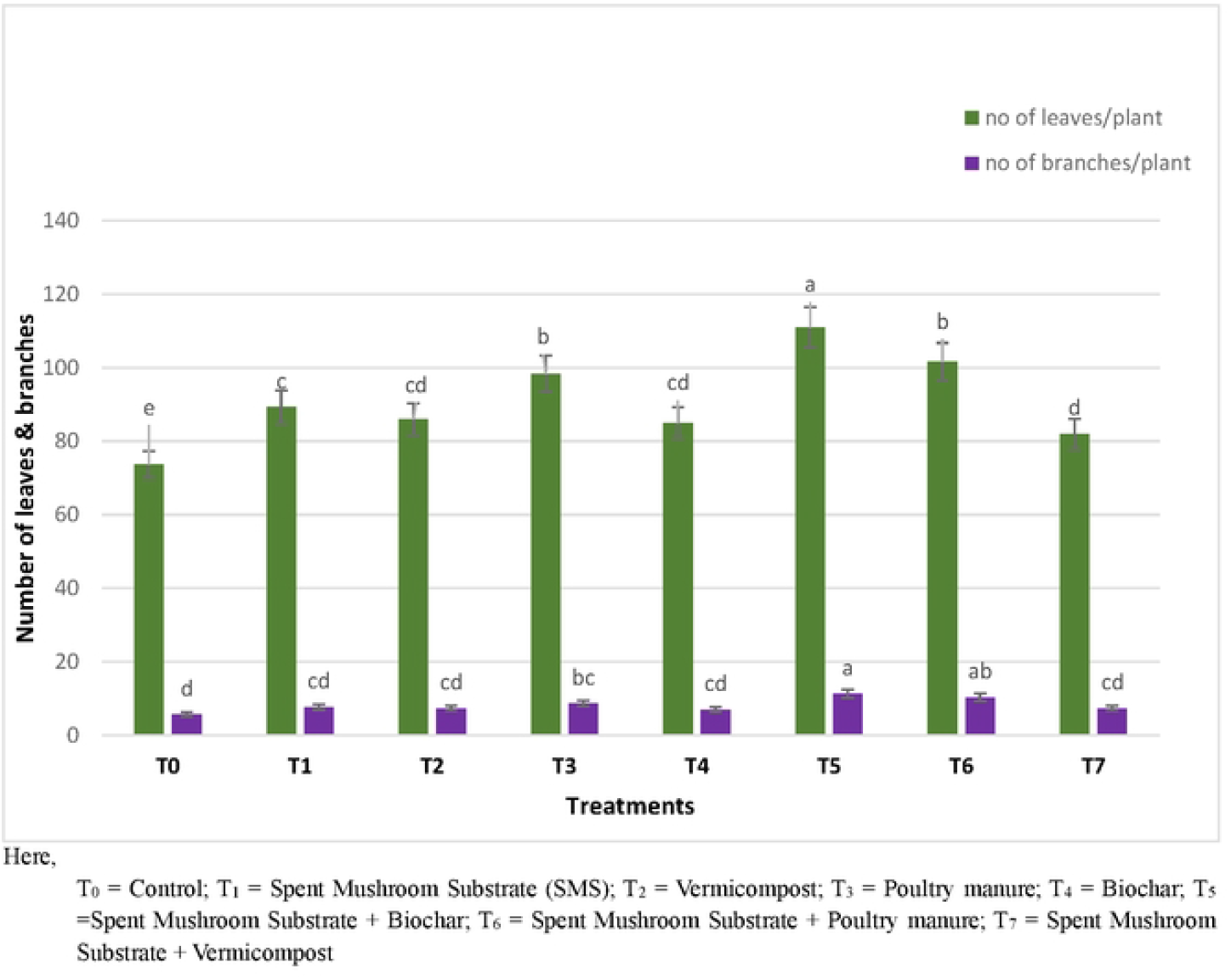
Impact of different treatments on the number of leaves and branches per plant of brinjal

### Study on yield and yield contributing attributes under different treatments

The effect of the selected treatments (T_0_-T_7_) against Fusarium wilt on yield and yield contributing characters of brinjal was studied and significant variation was observed. The treatments not only suppressed disease incidence but also positively impacted yield-contributing attributes. T_5_ demonstrated superior performance across all parameters, including the number of fruits per plant, number of fruits per plot, individual fruit weight, and fruit length. These metrics were significantly higher compared to the untreated control (T_0_), with T_5_ yielding 12.27 fruits per plant, 73.66 fruits per plot, an average fruit weight of 92.52 g, and a fruit length of 9.01 cm (Table 2).

**Table 2.** Impact of different treatments on yield contributing characters.

| Treatments | Number of fruit/plant | Number of fruit/plot | Individual weight of fruit (gm) | Fruit length (cm) |
| --- | --- | --- | --- | --- |
| T<sub>0</sub> | 3.77 d | 22.66 d | 73.32 bc | 7.84 b |
| T<sub>1</sub> | 4.77 cd | 28.66 cd | 73.87 bc | 8.56 ab |
| T<sub>2</sub> | 7.77 b | 46.66 b | 66.69 c | 8.28 ab |
| T<sub>3</sub> | 6.77 bc | 40.66 bc | 73.14 bc | 7.80 b |
| T<sub>4</sub> | 8.38 b | 50.33 b | 81.73 b | 8.81 ab |
| T<sub>5</sub> | 12.27 a | 73.66 a | 92.52 a | 9.01 a |
| T<sub>6</sub> | 8.55 b | 51.33 b | 92.09 a | 9.11 a |
| T<sub>7</sub> | 4.72 cd | 28.33 cd | 76.44 bc | 8.21 ab |
| CV (%) | 19.35 | 19.33 | 7.12 | 7.16 |

In terms of overall yield, T_5_ produced the highest values (865 g/plant, 5.19 kg/plot, and 12.71 tons/ha), outperforming all other treatments, including the untreated control (T_0_), which registered the lowest yield (277.67 g/plant, 1.66 kg/plot, and 4.08 tons/ha). T_6_ also significantly improved, ranking second across all yield parameters (Table 3). These findings underscore the effectiveness of spent mushroom substrate combined with biochar (T_5_) and poultry manure (T_6_) in enhancing growth, reducing disease incidence, and improving yield in brinjal cultivation.

**Table 3.** Impact of the selected treatments on yield of brinjal against Fusarium wilt.

| Treatments | Yield (g/plant) | Yield (kg/plot) | Yield (ton/ha) |
| --- | --- | --- | --- |
| T<sub>0</sub> | 277.67 d | 1.66 d | 4.08 d |
| T<sub>1</sub> | 353.67 cd | 2.12 cd | 5.19 cd |
| T<sub>2</sub> | 599.33 b | 3.59 b | 8.79 b |
| T<sub>3</sub> | 542.00 bc | 3.25 bc | 7.95 bc |
| T<sub>4</sub> | 691.00 ab | 4.14 ab | 10.12 ab |
| T<sub>5</sub> | 865.00 a | 5.19 a | 12.71 a |
| T<sub>6</sub> | 684.00 ab | 4.10 ab | 10.05 ab |
| T<sub>7</sub> | 364.67 cd | 2.18 cd | 5.33 |
| CV (%) | 21.97 | 21.97 | 22.17 |

## Discussion

### Pathogen identification

The identification of *Fusarium oxysporum* f. sp. *melongenae* as the causal agent of wilt in brinjal aligns with previous findings by [8, 16–17], where similar morphological characteristics and pathogenicity were observed. Consistent with these studies, the pathogen displayed whitish to light pink mycelium and the development of both 2 or 3 celled, slightly curved macroconidia and single-celled microconidia, confirming its identity through symptomatology and Koch’s postulates.

### Disease incidence

The study demonstrated that treatments with Spent Mushroom Substrate (SMS) in combination with biochar (T_5_) and poultry manure (T_6_) significantly reduced disease incidence compared to the untreated control (T_0_). The antifungal properties of SMS are well-documented, containing beneficial microorganisms like Trichoderma that exhibit antagonistic effects on soil-borne pathogens [18–20]. Furthermore, the integration of biochar in T_5_ enhanced disease suppression by improving soil health and inducing systemic resistance in plants [21–22]. In 2020, Singh and Kumar reported that up to a 50% reduction in *Fusarium* growth due to biochar’s inhibitory effects on chlamydospore development [23].

The combined application of SMS and poultry manure (T_6_) also showed significant disease reduction due to the presence of organic amendments like poultry manure that suppress the viability of soil-borne plant diseases [24]. These outcomes underscore the potential of bio-organic treatments as eco-friendly alternatives to chemical control methods for managing Fusarium wilt in brinjal.

### Growth parameters, yield and yield contributing attributes

The integration of SMS with biochar (T_5_) and poultry manure (T_6_) not only controlled disease incidence but also enhanced plant growth parameters and yield. The observed increases in plant height, leaf count, branch number, and overall yield were indicative of improved soil fertility and microbial activity. This is consistent with the findings that demonstrated that SMS applications enrich soil biodiversity and enhance crop productivity [25]. The authors reported that the application of SMS increased the diversity of fungi, including Tremellomycetes and Pezizomycetes for the SMS additive, while the levels of Mortierellomycetes, Pezizomycetes, and Leotiomycetes increased after the addition of poultry manure. The results suggest that incorporating SMS with other organic amendments could further improve soil properties and crop yields, supporting sustainable agriculture and reduced dependency on chemical inputs.

## Conclusion

The findings of this study illustrate that the application of Spent Mushroom Substrate (SMS) in combination with biochar (T_5_) and poultry manure (T_6_) significantly reduces the incidence of Fusarium wilt in brinjal while simultaneously enhancing plant growth parameters and yield. The bio-organic treatments not only suppress the pathogen effectively but also enrich soil health by promoting microbial diversity and improving soil structure. Therefore, the use of SMS-based organic amendments presents a sustainable and eco-friendly alternative to conventional chemical methods for managing soil-borne diseases in brinjal cultivation, contributing to improved crop productivity and sustainable agriculture.

## Acknowledgments

The author would like to express her heartfelt gratitude and appreciation to the Ministry of Science and Technology, Government of the People’s Republic of Bangladesh, for bestowing the prestigious National Science and Technology (NST) Fellowship upon the author. This esteemed award has played a crucial role in supporting and enabling the successful completion of this study.

## References

1. BBS. Bangladesh Bureau Statistics. Year Book of Agricultural Statistics, Statistics and Informatics Division, Ministry of Planning, Government of the People’s Republic of Bangladesh. 2023. Available from: https://bbs.gov.bd/site/page/3e838eb6-30a2-4709-be85-40484b0c16c6/Yearbook-of-Agricultural-Statistics

2. Naeem MY, Ugar S. Nutritional Content and Health Benefits of Eggplant. Turkish J. Agric. - Food Sci. Tech. 2019;7: 31–36.

3. Zeeshan, Ahmad M, Khan I, Shah B, Naeem A, Khan N, et al. Study on the management of Ralstonia solanacearum (Smith) with spent mushroom compost. J. Entomol. Zool. Studies. 2016;4(3): 114–121.

4. Sahoo R. Biorational Approach: An Alternative Approach to Control the Wilt Diseases of Crops. Acta Sci. Agric. 2022;6(9): 71–77.

5. Adhikary M, Begum, H, Meah M. Possibility Of Recovering Fusarium Wilt Affected Eggplants by Trichoderma. Int. J. Agril. Res. Innov. & Tech. 2017;7(1): 38–42.

6. Basco M, Bisen K, Keswani C, Singh HB. Biological management of Fusarium wilt of tomato using biofortified vermicompost. Mycosphere. 2017;8(3): 467–483.

7. Verma D, Didwana VS, Maurya B. Spent mushroom substrate: a potential sustainable substrate for agriculture. Int. J. Grid Distributed Comput. 2020;13(2): 104–109.

8. Adedeji, K. and Aduramigba, M. A. (2016). In vitro evaluation of spent mushroom compost on growth of Fusarium oxysporium f. sp lycopersici. Adv. Plants Agric. Res. 4(4): 332‒339.

9. Gudeta K, Bhagat A, Julka JM, Sinha R, Verma R, Kumar A, et al. Vermicompost and Its Derivatives against Phytopathogenic Fungi in the Soil: A Review. Horticulturae. 2022;8(311).

10. Gandhi A, Sundari US. Effect of Vermicompost Prepared from Aquatic Weeds on Growth and Yield of Eggplant (Solanum melongena L.). J Biofertil Biopesticide. 2012;3(5).

11. Richa, Kumar V, Singh J, Sharma N. Poultry Manure and Poultry Waste Management: A Review. Int.J.Curr.Microbiol.App.Sci. 2020;9(6): 3483–3495.

12. Jat MK, Ahir R, Choudhary S, Kakaraliya G. Management of coriander wilt (Fusarium oxysporum) through cultural practices as organic amendments and date of sowing. J. Pharmacog. Phytochem. 2017;6(5): 31–33.

13. Akanmu AO, Sobowale AA, Abiala MA, Olawuyi OJ, Odebode AC. Efficacy of biochar in the management of Fusarium verticillioides Sacc. Causing ear rot in Zea mays L. Biotechnology Reports 26. 2020.

14. Nutter FW, Esker PD, Coelho-Netto RA. Disease assessment concepts and the advancement made in improving the accuracy and precision of plant disease data. European J. Pl. Pathol. 2006;115: 99–103. doi: 10.1007/1-4020-5020-8_7

15. Das SN. Field efficacy of brinjal wilt with potential fungicides, biocontrol agents, and plant extracts. Nat. Volatiles & Essent. Oils. 2021;8(5): 9328–9341.

16. Altinok HH. (2005). First report of fusarium wilt of eggplant caused by Fusarium oxysporum f. sp. melongenae in Turkey. Plant Pathology. 2005;54: p 577.

17. Hassan HA. Biology and Integrated Control of Tomato Wilt Caused by Fusarium oxysporum lycopersici: A Comprehensive Review under the Light of Recent Advancement. J Bot Res. 2020;3(1): 84–99.

18. Ocimati W, Were E, Tazuba A F, Dita M, Zheng SJ. Spent Pleurotus ostreatus Substrate Has Potential for Managing Fusarium Wilt of Banana. J. fungi. 2021;7 (11): 946.

19. Yusidah I, Istifadah N. The abilities of spent mushroom substrate to suppress basal rot disease (Fusarium oxysporum f.sp cepae) in shallot. Int. J. Biosci. 2018;13 (1): 440– 448.

20. Salim HA, Simon S. Effect of carbendazim and solarized soil with Pseudomonas fluorescens, spent mushroom compost against Fusarium oxysporum f. sp. lycopersici in Tomato. European academic research. 2015;2(12). 15997–16010.

21. Medeiros EVD, Silva Lfd, Silva Jsad, Costa DPD, Souza Cafd, Berger LRR, et al. Biochar and Trichoderma spp. in management of plant diseases caused by soilborne fungal pathogens: a review and perspective. Research, Society and Development. 2021;10(15).

22. Poveda J, Gomez AM, Fenoll C, Escobar C. Use of biochar for plant pathogen control. The American phytopathological society. 2021;111: 1490–1499.

23. Singh N, Kumar A. (2020). Plant Disease Management through Bio-Char: A Review. Int. J. Curr. Microbiol. App. Sci. 2020;11: 3499–3510.

24. Zheng J, Wang L, Hou W, Han Y. (2022). Fusarium oxysporum Associated with Fusarium Wilt on Pennisetum sinese in China. Pathogens. 2022;11(999): 1–8.

25. Frac M, Pertile G, Panek J, Gryta A, Oszust K, Lipiec J, Usowicz B. Mycobiome Composition and Diversity under the Long-Term Application of Spent Mushroom Substrate and Chicken Manure. MDPI. 2021;11(3), 410.

